# TLR-mediated activation of synovial fibroblasts from osteoarthritis patients promotes chondrocyte dysfunction

**DOI:** 10.64898/2026.09.04.749398

**Authors:** Yujie Dai, Ao Gong, Nayar Durán-Hernández, Xu Liang, Xiaohui Liu, Linden J. Gearing, Pawel Durek, Yonghai Li, Stephan Oehme, Peihua Wu, Frederik F. Heinrich, Katrin Lehmann, Shilei Wu, Luis Lauterbach, Lorenz Pichler, Tazio Maleitzke, Stefanie Donner, Carsten Perka, Eicke Latz, Gerhard Krönke, Tobias Winkler, Mir-Farzin Mashreghi, Max Löhning, Ping Shen

## Abstract

**Objective:** Toll-like receptor (TLR) activation by cartilage-derived damage-associated molecular patterns contributes to osteoarthritis (OA) pathogenesis, but the role of synovial fibroblasts in this process remains incompletely understood. We investigated the TLR responsiveness of human OA synovial fibroblasts and determined how TLR activation influences fibroblast-mediated regulation of chondrocyte function.

**Design:** Primary synovial fibroblasts isolated from OA patients were characterized for TLR expression and stimulated with agonists targeting TLR1/2–TLR9. Inflammatory mediator production, matrix metalloproteinase expression, mitochondrial respiration, and transcriptomic responses were assessed. Functional effects on cartilage homeostasis were evaluated using autologous chondrocyte spheroid co-cultures.

**Results:** Approximately 50% of cells within OA synovial membranes were fibroblasts (CD45^-^CD31^-^PDPN^+^). Synovial fibroblasts expressed multiple TLRs at both the transcript and protein levels. Activation of TLR1/2, TLR4, TLR5, and TLR2/6 induced robust expression of IL-6, IL-8, G-CSF, MMP3, and MMP10, with minimal effects on mitochondrial respiratory function. Transcriptomic analysis revealed activation of inflammatory pathways together with enrichment of antigen presentation and protein translation programs. In autologous co-culture, TLR1/2-activated synovial fibroblasts promoted inflammatory and catabolic gene expression, suppressed anabolic gene expression, and impaired chondrocyte spheroid growth.

**Conclusions:** Human OA synovial fibroblasts are highly responsive to TLR activation and acquire a pro-inflammatory, cartilage-degrading phenotype that directly impairs chondrocyte homeostasis. These findings identify synovial fibroblasts as key effectors of innate immune signalling within the OA joint and support targeting shared TLR-mediated pathways in joint-resident cells as a potential disease-modifying therapeutic strategy.

## Introduction

Osteoarthritis (OA) is the most prevalent and disabling joint disease worldwide, affecting an estimated 7.6% of the global population (∼595 million individuals), a number projected to double by 2050 due to aging and rising obesity [1, 2]. Despite extensive research, no disease-modifying OA therapy has yet been approved, largely because the mechanisms driving OA onset and progression remain poorly understood.

Cartilage degradation is the hallmark of OA [3]. Cartilage integrity relies on a balance between anabolic and catabolic processes, supported by lubricants and nutrients supplied by the synovial membrane [4]. During OA progression related to aging or trauma, catabolic activity is amplified by mechanical stress and inflammation, leading to extracellular matrix breakdown, reduced mitochondrial respiratory capacity [5], dysregulated reactive oxygen species (ROS) production [6], and elevated inflammatory signalling [7]. As cartilage degrades, pro-inflammatory degradation products accumulate in the synovial microenvironment. Several of these products act as endogenous “damage signals”, activating Toll-like receptors (TLR) on macrophages and chondrocytes [5, 8]. Notably, 29 kDa fibronectin fragments and the aggrecan-derived 32-mer peptide function as TLR2 agonists [5, 9], and *Tlr2*-deficient mice develop less severe OA [10], supporting a pathogenic role for TLR signalling in OA. We previously showed that human chondrocytes express multiple TLRs, with TLR1/2 and TLR2/6 stimulation most potently suppressing chondrocyte spheroid growth by inhibiting extracellular matrix synthesis, promoting extracellular matrix degradation, inducing inflammatory cytokines, and impairing mitochondrial respiration [5].

OA is increasingly recognized as a whole-joint disease, affecting not only cartilage but also the synovial membrane [7, 11, 12]. Synovial fibroblasts are essential for joint homeostasis, producing synovial fluid [13] and extracellular matrix components [14]. In OA, these fibroblasts become dysregulated, secreting inflammatory mediators that recruit immune cells and cartilage-degrading enzymes that directly accelerate cartilage destruction [15]. Podoplanin (PDPN), a cell-surface glycoprotein, is highly expressed on synovial fibroblasts in inflamed joints and serves as a marker of activated fibroblast populations [16, 17]. PDPN⁺ fibroblasts comprise two major subsets: CD90⁺ sublining fibroblasts, which predominantly secrete inflammatory cytokines such as IL-6 and IL-8, and CD90⁻ lining fibroblasts, which preferentially produce cartilage-degrading enzymes including MMP1, MMP3, and ADAMTS4 [18]. CD34 and CD90 have been further reported to define PDPN^+^ fibroblast subpopulations with distinct functional and transcriptional profiles. CD34⁺ and CD34⁻CD90⁺ fibroblasts exhibit higher proliferative capacity and enhanced invasive and migratory abilities compared with CD34⁻CD90⁻ cells. In addition, CD34⁺ fibroblasts express higher levels of pro-inflammatory cytokine genes, including *IL6*, *CXCL12*, and *CCL2*, than CD34⁻ fibroblasts. In contrast, CD34⁻CD90⁻ fibroblasts show increased expression of genes associated with matrix remodeling and lubrication, such as MMP1, MMP3, and proteoglycan 4 (PRG4) [19]. Furthermore, CD34⁺CD90⁺ fibroblasts display elevated expression of osteogenic and chondrogenic differentiation markers, including runt-related transcription factor 2 (RUNX2) and aggrecan (ACAN) [20]. These findings highlight that cartilage degradation in OA arises from both chondrocyte dysfunction and synovial fibroblast–driven inflammation.

Fibroblasts from RA patients express TLR1–6, while TLR7–9 are generally undetectable, and stimulation with TLR ligands induces robust upregulation of MMPs [21]. As previous studies have used OA synovial fibroblasts only as a control group for RA fibroblasts [21–24], a comprehensive understanding of TLR expression and function in OA synovial fibroblasts is lacking. Hence, we here profiled the expression of TLR family members in synovial fibroblasts derived from OA patients and studied the responses of these cells to TLR stimulation with respect to possible influences on cartilage extracellular matrix homeostasis, inflammation, and energy metabolism. We show that TLR1/2, TLR4, TLR5, and TLR2/6 selectively induced the expression of inflammatory cytokines and cartilage-degrading enzymes, while exerting minimal effects on mitochondrial respiratory capacity of OA fibroblasts. Furthermore, using a trans-well co-culture system, we found that TLR1/2-prestimulated fibroblasts enhanced inflammatory and cartilage-catabolic gene expression while suppressing cartilage-anabolic gene expression in chondrocytes, resulting in impaired chondrocyte spheroid growth.

## Materials and Methods

### Patient samples

Synovial membranes, synovial fluid, and cartilage tissue were collected from OA patients undergoing knee arthroplasty. A total of 74 OA patients (39 women and 35 men; mean age: 69.7 ± 9.5 years; Table 1), all with clinically and radiographic image confirmed OA, provided informed consent for the use of clinical data and samples for research purposes. The study was approved by the responsible ethics committee, Ethikkommission der Charité – Universitätsmedizin Berlin (EA4/022/21).

**Table 1.** Demographic data of OA donors.

|  | Osteoarthritis patients |  |
| --- | --- | --- |
| | Number | Mean age ± SD (years) |
| Total | 74 | 69.67 ± 9.52 |
| Women | 39 | 70.73 ± 9.90 |
| Men | 35 | 68.49 ± 9.07 |

### Human synovial membrane cell and synovial fibroblast isolation

Synovial membrane samples were washed three times with 1% BSA and 5 mM EDTA in PBS to remove residual blood. Following excision of fat tissue, the synovial membrane was weighed and finely minced with scissors. The tissue fragments were then digested in a solution containing Type IV collagenase (4 mg/ml; Cell Signaling Technology) and DNase I (100 µg/ml; Roche) in DMEM supplemented with 1% penicillin-streptomycin. Digestion was carried out with shaking at 37°C for 60 minutes, after which the solution was filtered through 70 µm MACS SmartStrainers (Miltenyi) to obtain a single-cell suspension. Cells were washed three times, then stained with PDPN, CD45, and propidium iodide (PI) to exclude dead cells. PDPN⁺CD45⁻ synovial fibroblasts were gated and sorted by FACS, with sorting purities routinely exceeding 98%.

### Chondrocyte isolation and spheroid culture

Cartilage samples were washed three times with DPBS (Gibco) and finely minced with scalpels. The tissue fragments were digested with collagenase II (1 mg/ml) for 16 hours. The resulting suspension was filtered through a 70 µm MACS SmartStrainer (Miltenyi) to obtain single chondrocytes, which were then washed and resuspended in culture medium. To generate spheroids, chondrocytes were resuspended in serum-free DMEM-High Glucose medium (Sigma Aldrich) supplemented with 0.1 µM dexamethasone (Sigma Aldrich; D2915), 40 µg/ml L-proline (Sigma Aldrich; P5607), 6.25 µg/ml insulin-transferrin-sodium selenite supplement (Sigma Aldrich; I1884), 0.1 mg/ml sodium pyruvate (AppliChem; A4859), 1.25 mg/ml bovine serum albumin (Sigma Aldrich; A9418), 1% penicillin-streptomycin (Gibco; 15140-122), 50 µg/ml 2-phospho-L-ascorbic acid trisodium salt (Sigma Aldrich; A8960), 5.35 µg/ml linoleic acid (Sigma Aldrich; L1012), and 10 ng/ml TGF-β1 (Peprotech; 100-21-5). For spheroid formation, 2.5 × 10⁵ cells were transferred into 15 ml Falcon tubes and centrifuged at 500 × g for 5 minutes. After carefully removing the supernatant, 500 µl of fresh spheroid medium was added without disturbing the cell pellet. Cultures were maintained in a hypoxic incubator (4% O₂, 5% CO₂, 37°C), and medium was renewed every three days using a hypoxic chamber workstation (BioSpherix X3, Xvivo system).

### Trans-well coculture

Isolated synovial fibroblasts were stimulated with or without the TLR2 agonist Pam3CSK4 (2 μg/ml) for 3.5 days in a 24-well plate. After stimulation, fibroblasts were harvested, washed twice, resuspended in chondrocyte spheroid culture medium (survival was comparable to standard fibroblast culture medium), counted, and seeded into a fresh 24-well plate at the same cell density. A 0.4 µm Trans-well insert (Millipore) was then placed into each well, and 0.5 ml of culture medium was added to the upper chamber. Chondrocyte spheroids derived from the same patients were transferred onto the inserts, allowing fibroblasts in the lower chamber and chondrocyte spheroids in the upper chamber to share medium while remaining physically separated. Cocultures were maintained in a hypoxic incubator (4% O₂, 5% CO₂, 37°C) with medium refreshed every 3.5 days. Supernatants were collected at each medium change and stored at –20°C for further analyses.

### TLR stimulation

Isolated synovial fibroblasts were stimulated with 2 µg/ml of Pam3CSK4, PolyI:C, LPS, Flagellin, Pam2CSK4, Imiquimod, ssRNA, or CpG (InvivoGen), which act as agonists of TLR1/2, 3, 4, 5, 2/6, 7, 8, and 9, respectively.

Detailed information for Synovial fluid cell isolation, Flow Cytometric Analysis, mRNA isolation and quantitative reverse transcription PCR, Synovial fibroblast TaqMan qPCR, RNA-sequencing analysis, Bio-plex analysis, Nitric oxide quantification by modified Griess reaction, MitoSpy, TMRM, MitoSox, and DCFDA staining, Mito Stress Test seahorse assay, Immunofluorescence Staining of Synovial Membrane, and Statistics, please see Supplementary Methods.

### Study approval

The study was approved by the Ethics Committee of Charité – Universitätsmedizin Berlin (EA4/022/21).

## Results

### Synovial fibroblasts from osteoarthritis patients exhibit broad expression of TLR family members

To optimize synovial membrane digestion, we tested multiple conditions using type IV collagenase (1, 2, or 4 mg/ml) or Liberase TL (100 or 200 µg/ml) combined with DNase I (0.1 mg/ml) on equal amounts of pre-minced synovial membrane tissue from the same patients (Figure 1A). Digestion with type IV collagenase (4 mg/ml) and DNase I (0.1 mg/ml) yielded the highest viable cell recovery and was thus used throughout the study. Single-cell suspensions were immediately analysed by flow cytometry to identify PDPN⁺ fibroblasts and immune cell populations (Figure 1B). Across OA synovial membrane samples, approximately 50% of cells were PDPN⁺CD45⁻ fibroblasts and about 28% were CD45⁺ immune cells, including approx. 20% CD14^+^ monocytes/ macrophages. On average, synovial tissue contained about 0.5 million fibroblasts and 0.3 million immune cells per gram. Thus, OA synovial membranes are dominated by PDPN⁺ fibroblasts with diverse immune cells, whereas synovial fluid contains only very few fibroblasts and its cellular fraction is primarily composed of immune cells (Supplementary Figure 1).

**Fig. 1.**
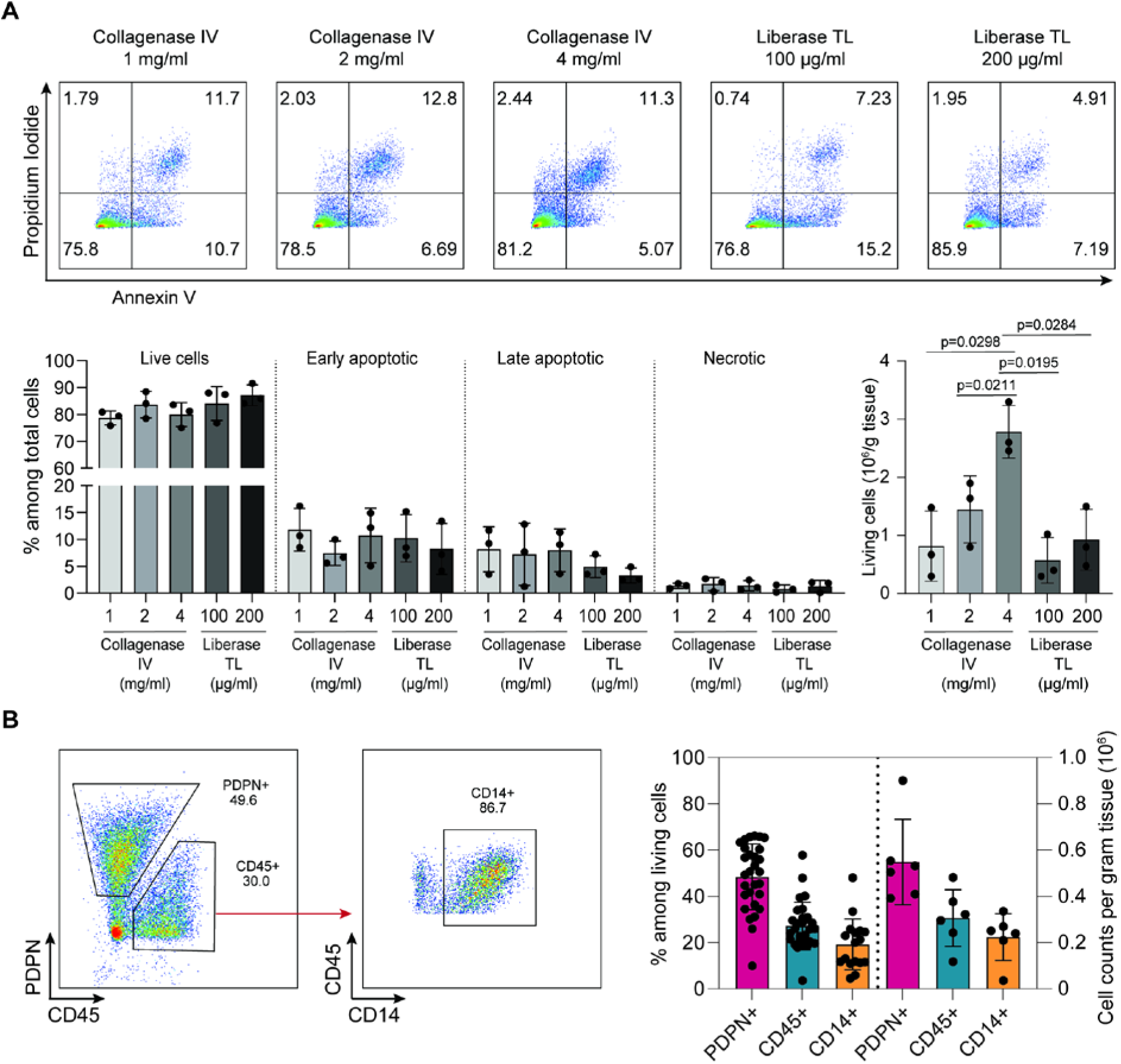
Optimized synovial cell isolation procedure revealed the composition of fibroblasts and immune celles in the synovail membrane of OA patients. Synovial membrane tissues were collected from OA patients immediately after surgery and were minced to small pieces. **(A)** Equal amounts of tissue were digested with collagenase IV (CIV) at concentrations of 1, 2, or 4 mg/ml or Liberase TL (TL) at concentrations of 100 µg/ml or 200 µg/ml combined with DNase I (100 µg/ml) for 1 hour in continuous oscillation (175 rpm) at 37°C. Single-cell suspensions were then stained with annexin V (AV) and propidium iodide (PI) to assess cell viability. Top, representative FACS plots. Bottom, quantification of live (AV^-^ PI^-^), early apoptotic (AV^+^PI^-^), late apoptotic (AV^+^PI^+^), and necrotic (AV^-^PI^+^) cells in percentages and cell numbers (n=3, mean ± SD). Data were compared using paired, one-tailed t-tests. **(B)** Synovial membrane was digested with collagenase IV and single-cell suspensions were analyzed by flow cytometry. Left, representative FACS plots. Right, quantification of fibroblasts (PDPN⁺CD45⁻), total immune cells (PDPN⁻CD45⁺), and monocytes/macrophages (CD45⁺CD14⁺), shown as percentages (n = 17, mean ± SD) and absolute cell numbers (n = 6, mean ± SD).

Synovial fibroblasts, particularly TLR-activated synovial fibroblasts, have been identified as key contributors to RA pathogenesis by promoting inflammation and joint destruction [25–27]. However, to date, few studies have directly and systemically examined TLR expression in OA synovial fibroblasts or their potential role in OA pathogenesis. Given the emerging recognition of low-grade innate inflammation in OA as a disease driver, it is essential to determine whether OA synovial fibroblasts express TLRs and, if so, which family members are preferentially expressed. To assess *ex vivo* the expression of TLRs and the downstream signaling molecules *MYD88* and *TRIF* in OA synovial fibroblasts, actinomycin D was used during cell isolation to inhibit *de novo* RNA synthesis. Live fibroblasts (PDPN⁺CD45⁻) were isolated by FACS with high purity (Supplementary Figure 2) and analyzed by TaqMan quantitative PCR (qPCR). We detected robust expression of *TLR1*, *TLR2*, *TLR3*, *TLR4*, *TLR5*, *TLR6*, and *TLR10* along with expression of the TLR signaling adaptor molecules *MYD88* and *TRIF*, whereas *TLR7*, *TLR8*, and *TLR9* were barely detectable (Figure 2A). Immunofluorescence analysis of freshly collected OA synovial membrane showed prominent TLR2 expression on PDPN⁺ fibroblasts (Figure 2B). While TLR1, TLR2, TLR4, TLR5, and TLR6 are traditionally classified as cell-surface receptors, multiple studies have shown that they also undergo intracellular trafficking and transient intracellular localization [28–31]. Therefore, to obtain a comprehensive view of TLR protein expression in synovial fibroblasts, we first conducted intracellular flow cytometric analysis to assess both surface and intracellular TLR expression in freshly isolated synovial membrane cells. This analysis revealed a uniform rightward shift of the entire cell population relative to the unstained control, rather than the emergence of a distinct TLR-positive subpopulation (Figure 2C), indicating that the TLR family members are expressed, albeit at different levels, across the whole PDPN⁺ fibroblast population. Overall, synovial PDPN⁺ fibroblasts displayed a broad repertoire of expressed TLRs, ranging from TLR1 to TLR9. Interestingly, although the transcripts for TLR7, TLR8, and TLR9 were minimally expressed in synovial fibroblasts, protein expression was still detectable, consistent with previous reports of uncoupling between TLR mRNA and protein expression in immune tissues [32, 33]. To further understand the surface presentation of TLRs (TLR1, TLR2, TLR4, TLR5, and TLR6), we performed cell surface staining for the same six samples used for intracellular TLR staining. Surface-only staining for TLR2 showed a distinct TLR2^high^ population, while the detection of the other four TLRs still displayed a shift in the TLR staining pattern (Figure 2D), albeit to a lesser extent, in comparison to their corresponding intracellular staining (Figure 2E). This is consistent with previous observations suggesting that membrane TLRs (TLR1, TLR2, TLR4, TLR5, and TLR6) can also remain localized intracellularly [30, 34–37].

**Fig. 2.**
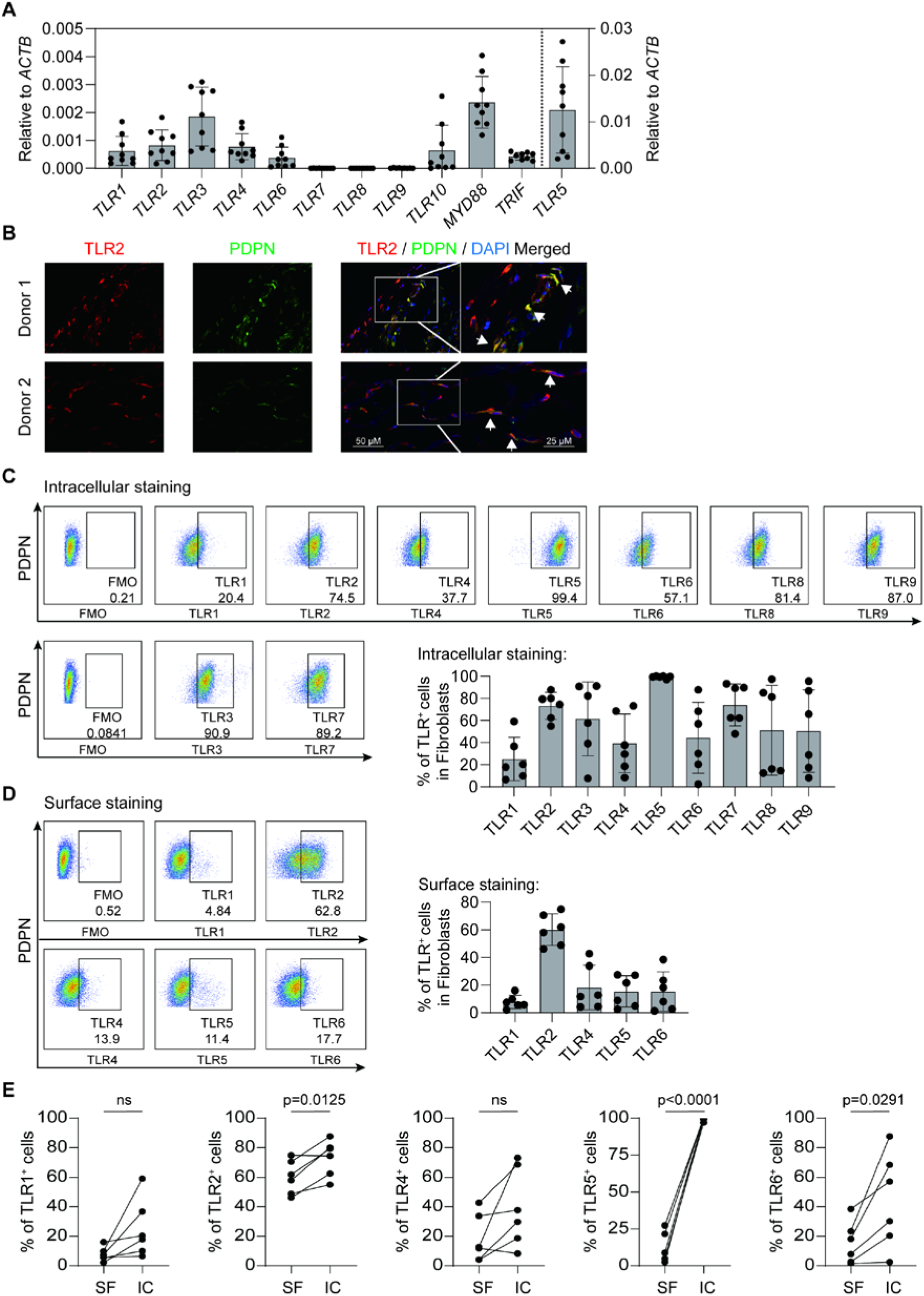
Synovial fibroblasts from OA patients exhibit broad expression of TLR family members. PDPN⁺CD45^−^ synovial fibroblasts were isolated by fluorescence-activated cell sorting (FACS) and immediately subjected to RNA extraction. (**A**) Expression of TLR family members, *MYD88*, and *TRIF* was detected by TaqMan quantitative PCR (n = 9, mean ± SD). (**B**) Synovial membrane tissues were processed for immunofluorescence staining of TLR2 and PDPN. (**C**) Freshly isolated synovial cells were stained with antibodies against PDPN, CD45, and CD14, followed by fixation, permeabilization, and intracellular staining for TLR1–TLR9. Top, representative FACS plots. Bottom, frequencies of TLR-expressing fibroblasts (PDPN⁺CD45⁻CD14⁻) (n = 6, mean ± SD). (**D**) Freshly isolated synovial cells were simultaneously stained with anti-PDPN, anti-CD45, anti-CD14, and antibodies against TLR1, TLR2, TLR4, TLR5, or TLR6. Left, representative FACS plots. Right, frequencies of surface TLR-expressing fibroblasts (PDPN⁺CD45⁻CD14⁻) (n = 6, mean ± SD). **(E)** Freshly isolated synovial cells were subjected to surface staining of PDPN, CD45, and CD14 to identify PDPN^+^CD45^-^CD14^-^ fibroblasts, as well as surface or intracellular staining of TLR1, TLR2, TLR4, TLR5, and TLR6 (n = 6). Data were compared using paired two-tailed t-test (ns, p > 0.05).

### Distinct subsets of synovial fibroblasts from OA patients express similar TLR2 levels

Synovial fibroblasts can be categorized into lining fibroblasts (CD90⁻) and sub-lining fibroblasts (CD90⁺) [38]. We compared the percentage of TLR2-expressing cells between these two fibroblast subsets using surface staining and found that TLR2 was similarly expressed by both lining and sub-lining fibroblasts (Figure 3A). Since CD34 and CD90 have been further reported to define fibroblast subpopulations with distinct functional and transcriptional profiles [20], we next compared the percentage of TLR2-expressing cells among the four fibroblast subsets. We found that TLR2 was similarly expressed by CD90⁻CD34⁺, CD90⁻CD34⁻, CD90⁺CD34⁻, and CD90⁺CD34⁺ fibroblasts (Figure 3B).

**Fig. 3.**
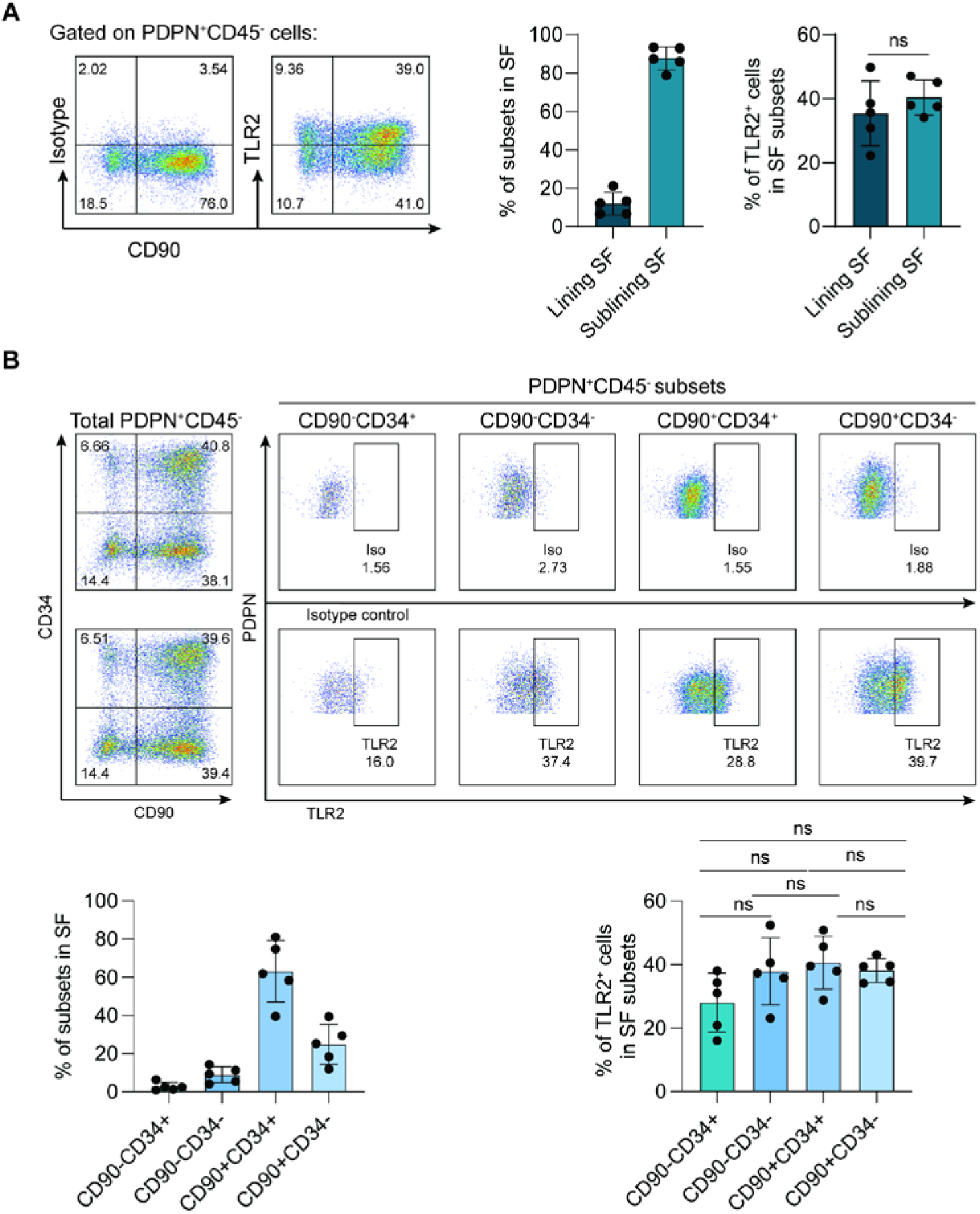
Distinct subsets of synovial fibroblasts from OA patients similarly express TLR2. Freshly isolated synovial membrane cells were simultaneously stained with anti-PDPN, anti-CD45, anti-CD14, anti-CD90, anti-CD34, and anti-TLR2 or its corresponding isotype control. (**A**) Left, representative FACS plots for the expression of TLR2 by lining (PDPN^+^CD90⁻) and sublining (PDPN⁻CD90^+^) synovial fibroblasts (SF). Right, quantification of TLR2-expressing fibroblast subsets (n = 5, mean ± SD). (**B**) Top, representative FACS plots for the expression of TLR2 by CD90⁻CD34⁺, CD90⁻CD34⁻, CD90⁺CD34⁺, and CD90⁺CD34⁻ synovial fibroblasts (SF). Bottom, quantification of TLR2-expressing cells in fibroblast subsets (n = 5, mean ± SD) Data were compared using one-way ANOVA (ns, p > 0.05).

### TLR activation drives inflammatory and cartilage-degrading responses in OA synovial fibroblasts

To evaluate the effects of TLR stimulation on OA synovial fibroblasts, total live PDPN⁺ fibroblasts were sorted by FACS to high purity (Supplementary Figure 2) without distinguishing fibroblast subsets, as these subsets express similar levels of TLRs. Cells were then stimulated with agonists for TLR1/2, TLR3, TLR4, TLR5, TLR2/6, TLR7, TLR8, or TLR9 (Supplementary Figure 3A). TLR stimulation did not affect fibroblast viability, as comparable levels of live cells were detected across all conditions, including unstimulated controls, after a 4-day culture (Supplementary Figure 3B).

To assess the transcriptional impact of TLR stimulation, RNA-sequencing was conducted for fibroblasts treated with each TLR agonist, as well as the unstimulated controls, following 4 days of culture. After correcting for donor effects, multidimensional scaling (MDS) analysis revealed that stimulation of TLR1/2, TLR4, TLR5, and TLR2/6 caused OA fibroblasts to cluster separately from unstimulated controls, whereas stimulation of TLR3, TLR7, TLR8, and TLR9 had minimal effects (Supplementary Figure 4A), indicating distinct response patterns between cell-surface and intracellular TLRs. Confident effect sizes (confects) plots identified genes that were significantly differentially regulated by each TLR stimulation (Figure 4A). Specifically, TLR1/2, TLR4, TLR5, and TLR2/6 stimulation resulted in significant upregulation of 111, 275, 31, and 185 genes, respectively, and downregulation of 12, 155, 9, and 33 genes, respectively. Although a number of gene expression changes were observed for TLR7, the magnitude of these changes was much weaker. In contrast, TLR3, TLR8, and TLR9 induced minimal gene expression changes (FDR < 0.05) (Figure 4B).

**Fig. 4.**
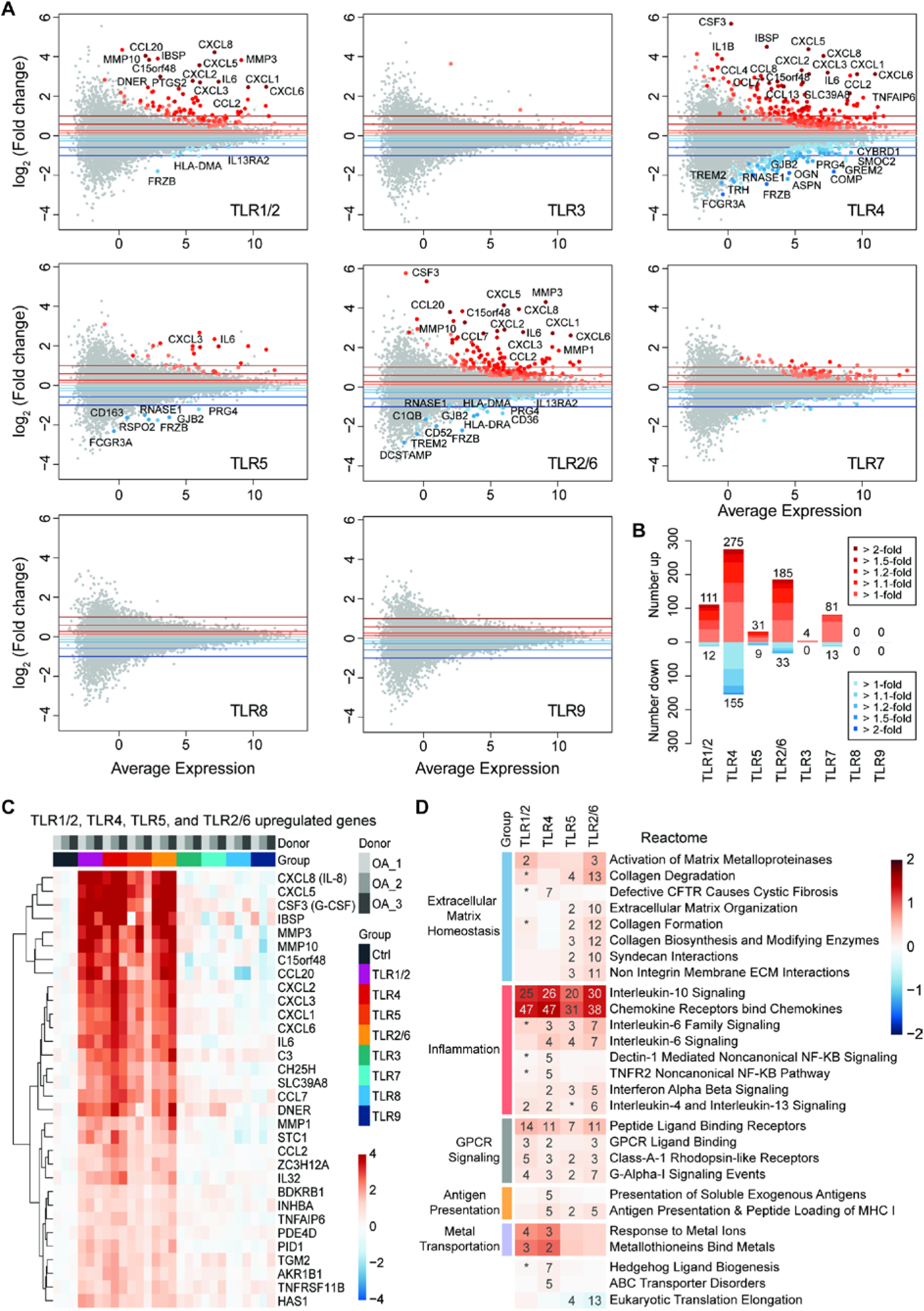
TLR activation drives inflammatory and cartilage-degrading responses in OA synovial fibroblasts. RNA-seq was performed on isolated synovial fibroblasts from three OA patients that were cultured with or without various types of TLR agonists for 4 days. (**A**) Mean-difference plot with confident effect sizes (confects) to identify genes that were significantly differentially regulated by each TLR stimulation. Selected genes are labelled. (**B**) Stacked bar plot shows the numbers of genes that met various confect thresholds. (**C**) Heat map displaying those genes that were commonly upregulated by TLR1/2, TLR4, TLR5, and TLR2/6 stimulation. Relative log_2_ gene expression compared to control, truncated at ±4. (**D**) Heat map combining the top ten gene sets from the Reactome gene set collection withTLR1/2, TLR4, TLR5, and TLR2/6 stimulation. Mean gene set log_2_ fold change, with significance indicated (−log_10_ FDR-adjusted p-value threshold).

As TLR1/2, TLR4, TLR5, and TLR2/6 signaling pathways share the adaptor protein MYD88 (with TLR4 also engaging TRIF), a shared transcriptional response was anticipated. Indeed, these stimulations upregulated inflammatory chemokines and cytokines—including *IL6*, *CXCL6*, *CXCL1*, *CXCL3*, *CXCL2*, *CCL20*, *CSF3* (G-CSF), *CXCL5*, and *CXCL8* (IL-8)—as well as the cartilage-degrading enzymes *MMP1*, *MMP3*, and *MMP10* (Figure 4C, and Supplementary Figure 4B). Pathway analysis using the Reactome database revealed that upregulated genes were primarily involved in extracellular matrix homeostasis, inflammation, GPCR (G Protein-Coupled Receptor) signaling, antigen presentation, metal transport, Hedgehog ligand biogenesis, and ATP-binding cassette (ABC) protein disorders, while downregulated genes were mainly associated with translation-related processes (Figure 4D). A similar pattern was observed in our previous study of TLR1/2-stimulated human chondrocytes [5]. Differential expression of IL-6, IL-8, and G-CSF (Figure 5A), as well as the cartilage-degrading enzymes *MMP1*, *MMP3*, and *MMP13* (Figure 5B), was further confirmed at the mRNA and/or protein level. These findings indicate that TLR stimulation enhances the inflammatory and cartilage-degrading activity of synovial fibroblasts, potentially promoting joint degeneration in OA patients. Overall, beyond the previously reported IL-6 induction by TLR2 stimulation (39), our data demonstrate a broader effect of TLR activation on the inflammatory phenotype of OA synovial fibroblasts, including upregulation of both inflammatory cytokines and cartilage-degrading enzymes.

**Fig. 5.**
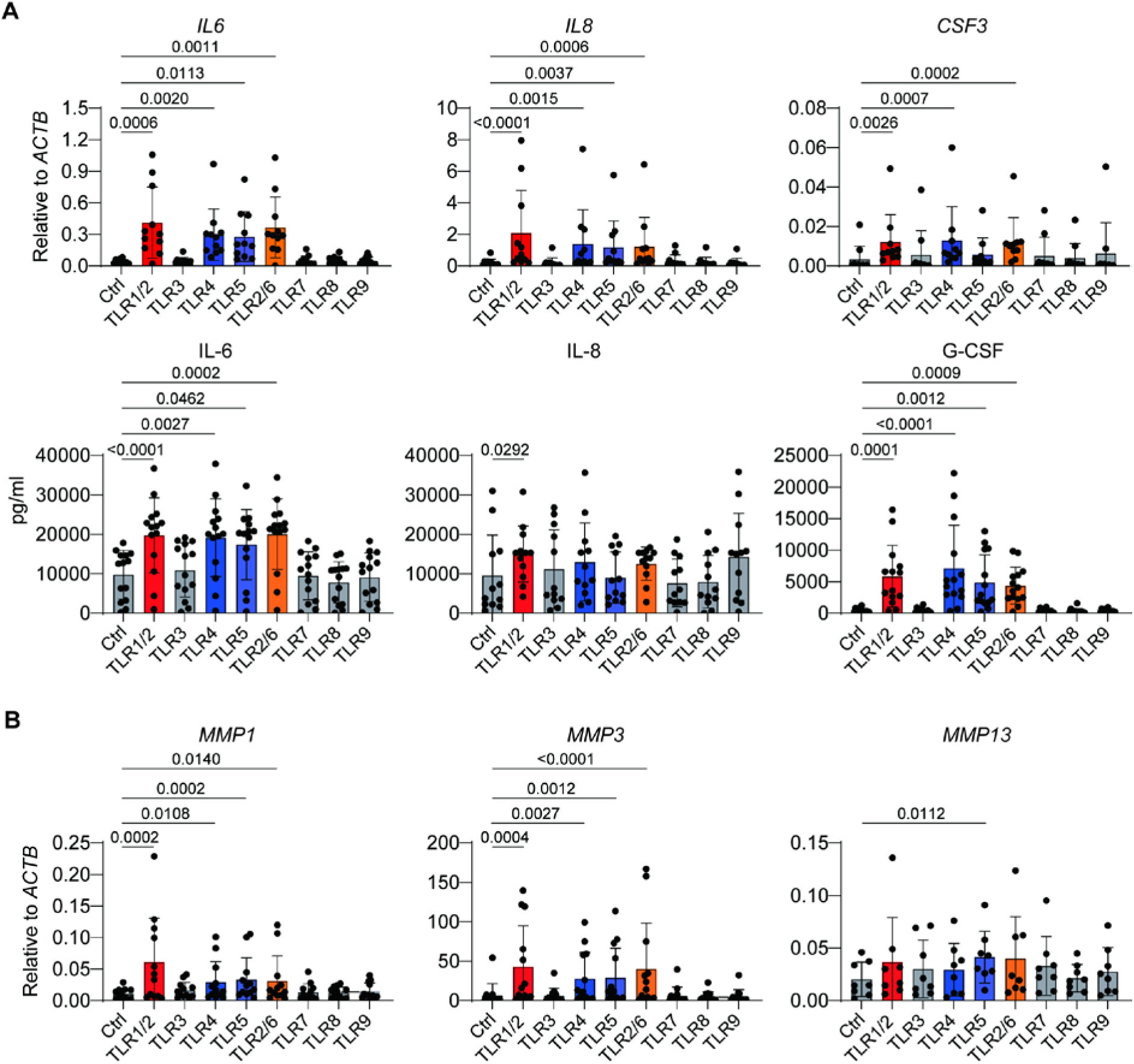
TLR activation drives inflammatory and cartilage-degrading responses in the synovial fibroblasts of OA patients. Freshly isolated human synovial fibroblasts were cultured with or without various types of TLR agonists for 4 days. (**A**) Expression of inflammatory cytokines IL-6, IL-8, and G-CSF was determined at mRNA level using qPCR (top; n=11, mean ± SD) and at protein level using Bio-Plex (bottom; n=12–14, mean ± SD). (**B**) Expression of cartilage-degrading enzymes was determined at mRNA level using qPCR (n=8– 12, mean ± SD). Data were compared using one-way ANOVA.

### TLR1/2 stimulation does not alter mitochondrial respiration or glycolytic capacity of OA synovial fibroblasts

Previously, we reported that TLR1/2 stimulation impairs mitochondrial respiration in human chondrocytes [5]. To investigate whether OA synovial fibroblasts are similarly affected, cells were stimulated with the TLR1/2 agonist Pam3CSK4 (P3C4) for four days. MitoSpy staining showed comparable mitochondrial mass between control and P3C4-stimulated fibroblasts, indicating that TLR1/2 activation does not alter mitochondrial content (Figure 6A). We then performed a Seahorse Mito Stress Test on day 3 of culture to assess mitochondrial respiration and glycolytic activity via oxygen consumption rate (OCR) and extracellular acidification rate (ECAR), respectively. Surprisingly, P3C4 stimulation had little effect on the metabolic activity of synovial fibroblasts, as both basal and maximal OCRs were similar to controls, and glycolytic activity remained unchanged (Figure 6B). Consistent with these findings, mitochondrial membrane potential, measured by TMRM staining, was comparable between control and P3C4-treated fibroblasts (Figure 6C). In contrast to RA synovial fibroblasts [39], OA fibroblasts did not upregulate NOS2 expression or nitric oxide (NO) production following TLR stimulation (Figure 6D). Given the established role of NO in impairing mitochondrial respiration in chondrocytes, this likely explains why TLR1/2 stimulation had little impact on mitochondrial function in OA synovial fibroblasts. Additionally, total intracellular reactive oxygen species (ROS) levels were unchanged, although mitochondrial ROS showed a slight but significant increase after TLR1/2 activation (Supplementary Figure 5).

**Fig. 6.**
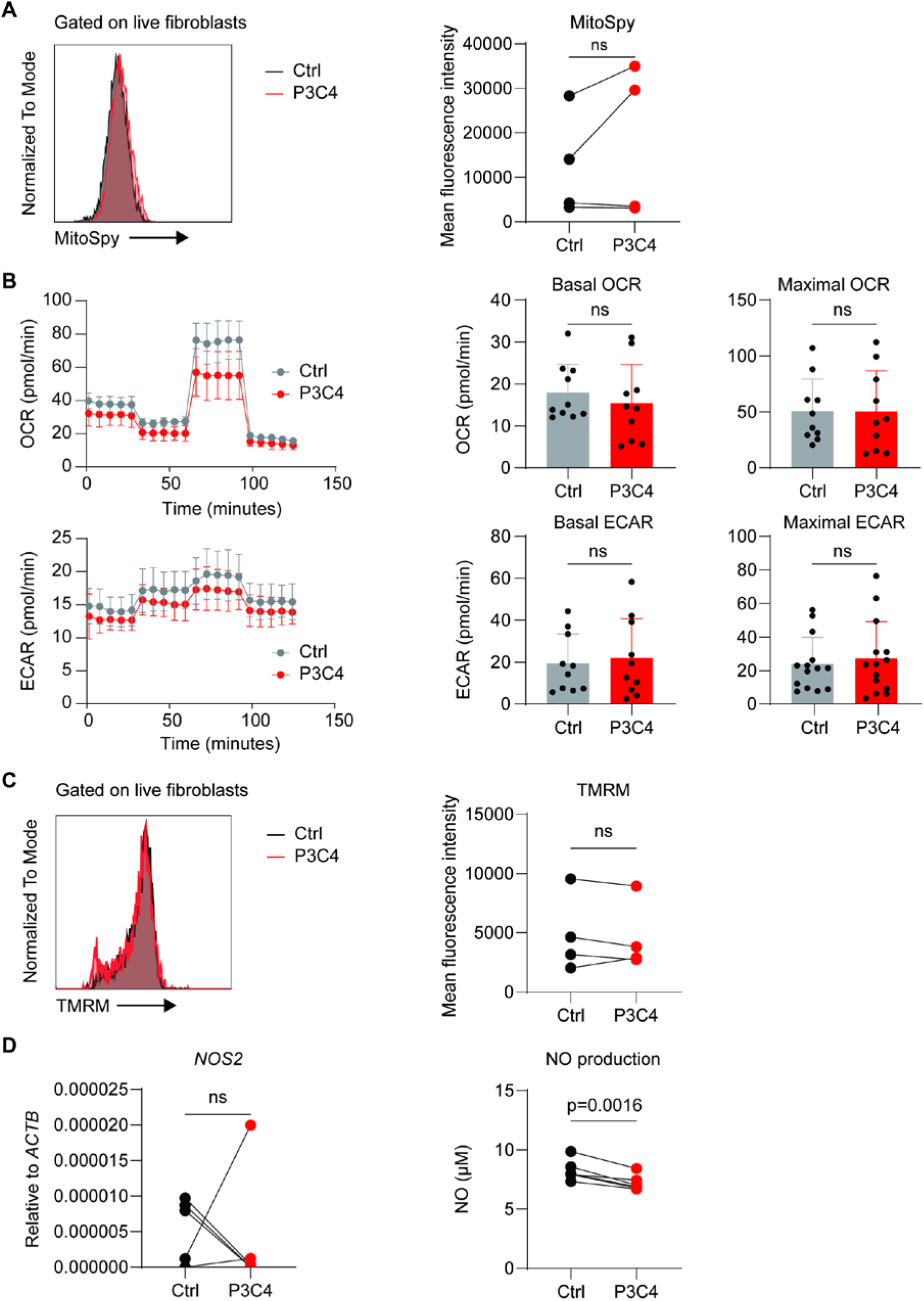
TLR activation does not alter their mitochondrial respiration or glycolytic capacity. Freshly purified synovial fibroblasts were stimulated with Pam3CSK4 (P3C4), the agonist of TLR1/2, or were left unstimulated as a control (Ctrl). Four days later, cells were harvested and subjected to MitoSpy staining to detect total mitochondrial mass (**A**, n=4), seahorse assays to determine mitochondrial respiration capacity and glycolytic activity using Mito Stress Test Kits (**B**, n = 10, mean ± SD). OCR: Oxygen Consumption Rate; ECAR: Extracellular Acidification Rate. Data were compared using paired two-tailed t test (ns, p > 0.05). cells were subjected to TMRM staining to detect mitochondrial membrane potential (**C**, n=4). *NOS2* mRNA expression in fibroblasts was quantified using qPCR (left), and NO production (right) in the supernatant was determined using Griess reaction (**D**, n=6).

These results indicate that, unlike chondrocytes, OA synovial fibroblasts maintain metabolic activity and ROS homeostasis despite robust TLR1/2-mediated inflammatory and cartilage-degrading responses.

### TLR1/2 pre-stimulation of synovial fibroblasts impairs chondrocyte function and inhibits chondrocyte spheroid growth

As TLR stimulation induces OA synovial fibroblasts to secrete inflammatory cytokines (IL-6, IL-8, G-CSF, IL-1β) and upregulate the expression of cartilage-degrading enzymes (*MMP1*, *MMP3*, *MMP10*, *MMP13*), we next investigated whether TLR-stimulated fibroblasts could impair chondrocyte function in a Trans-well co-culture system (Figure 7A). Cultures were maintained under 4% O₂. Three days later, chondrocyte viability was assessed (Supplementary Figure 6), and expression of inflammatory factors was quantified. While synovial fibroblasts (SF) did not alter the mRNA expression of *IL6*, *CXCL8*, or *CSF3*, TLR1/2-prestimulated synovial fibroblasts (P3C4-SF) markedly increased the expression of the inflammatory factors IL-6, CXCL8 (IL-8), and G-CSF in chondrocytes at both the mRNA and protein levels (Figure 7B). With regard to the expression of chondrocyte anabolic and catabolic factors, even without pre-stimulation, SF reduced the expression of the cartilage-anabolic factor *COL2A 1*(Figure 8A)., demonstrating the anti-anabolic impact of OA fibroblasts on cartilage tissue. Notably, P3C4-SF fibroblasts suppressed *COL2A1* expression even more significantly. For cartilage-catabolic factors, unstimulated fibroblasts tended to modestly increase *MMP3* and *MMP13* expression, whereas TLR1/2-prestimulated fibroblasts significantly elevated the expression of these cartilage-degrading enzymes in chondrocytes. We then assessed the impact of TLR1/2-prestimulated synovial fibroblasts on chondrocyte spheroid growth. Using the same co-culture system, fibroblasts and chondrocyte spheroids were maintained together for 28 days, and spheroid weight was measured (Figure 8B). TLR1/2-prestimulated fibroblasts suppressed spheroid growth most potently, resulting in more than 60% reduction in weight. Taken together, these results demonstrate that TLR1/2 pre-stimulation drives OA synovial fibroblasts to further promote dysfunction of chondrocytes from the same patients.

**Fig. 7.**
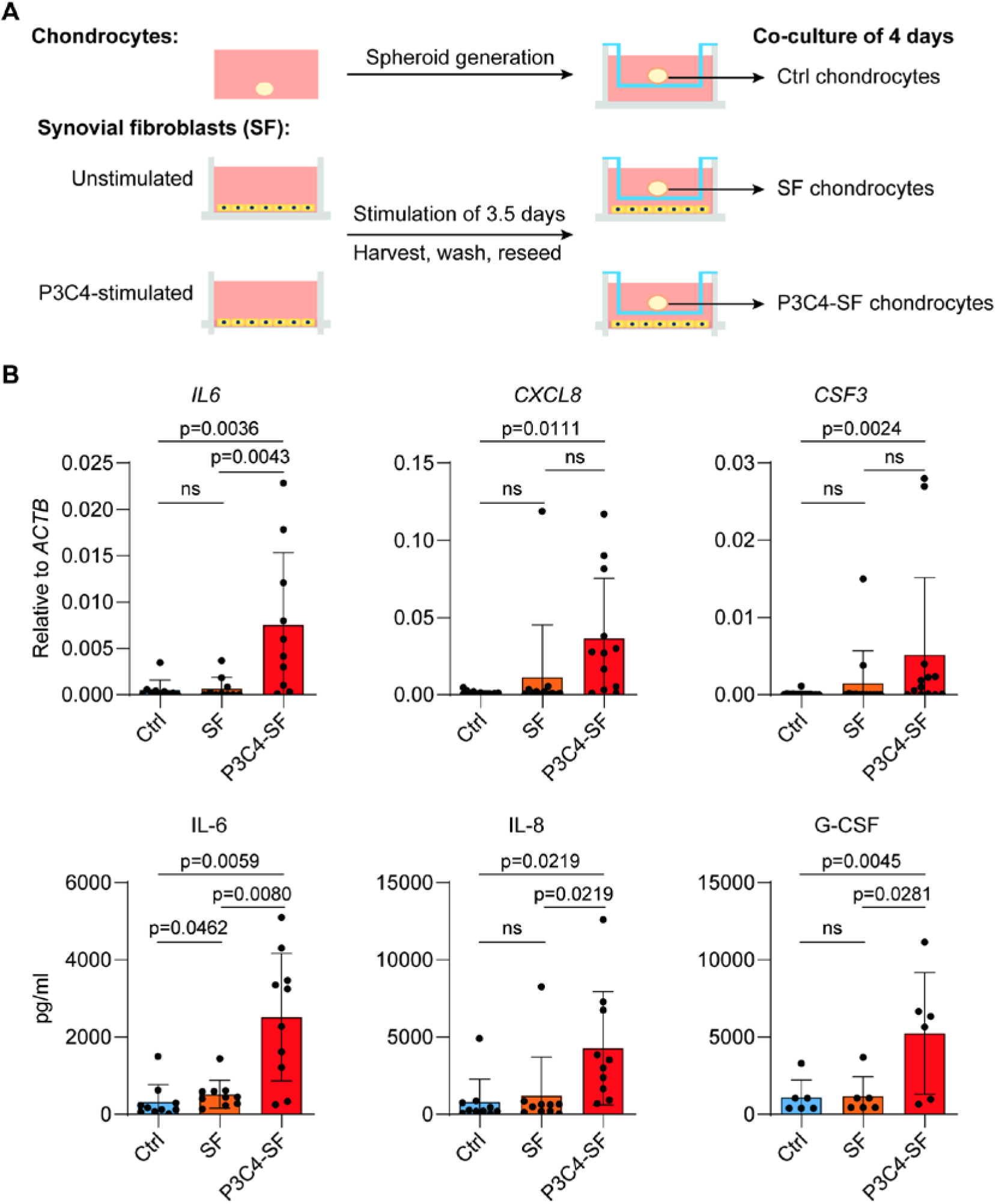
TLR1/2 pre-stimulation of synovial fibroblasts induces inflammatory features in chondrocyte spheroid. **(A)** Setup of Trans-well coculture system. Both chondrocytes and synovial fibroblasts (SF) were isolated from the same patient. While chondrocytes were subjected to spheroid generation, synovial fibroblasts were stimulated with or without the TLR2 agonist Pam3CSK4 for 3.5 days. After stimulation, fibroblasts were harvested, washed, and reseeded into a fresh 24-well plate at the same cell density. A 0.4 µm Transwell insert (Millipore) was then placed into each well and chondrocyte spheroids derived from the same patient were placed onto the inserts, allowing fibroblasts in the lower chamber and chondrocyte spheroids in the upper chamber to share a growth environment while remaining physically separated. (**B**) After a 4-day co-culture, *IL6* (n=10), *CXCL8* (n=12), and *CSF3* (n=13) mRNA expression in chondrocytes was assessed using qPCR (top) and IL-6 (n=10), IL-8 (n=10), and G-CSF (n=6) secretion in the supernatant was determined using Bio-Plex assay (bottom).

**Fig. 8.**
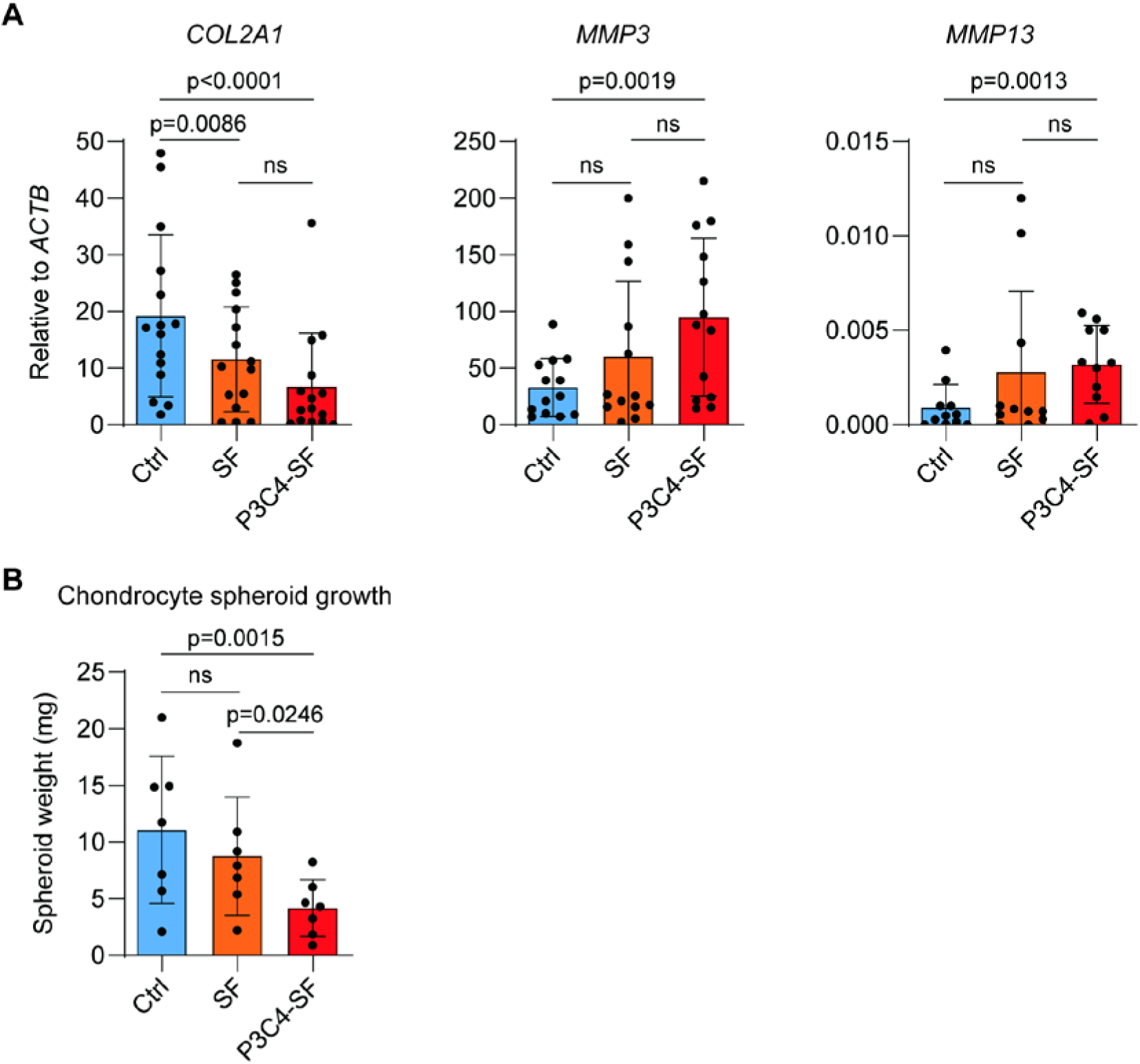
TLR1/2 pre-stimulation of synovial fibroblasts impairs inhibits chondrocyte spheroid growth. (**A**) After a 4-day co-culture, mRNA expression of *COL2A1* (n=15), *MMP3* (n=13), and *MMP13* (n=11) in chondrocytes was assessed by qPCR. (**B**) Chondrocyte spheroids and fibroblasts were cultured for 28 d. The weight of each spheroid was determined (n = 7, mean ± SD). Data were compared using one-way ANOVA (ns, p > 0.05).

## Discussion

We determined the cellular composition of the synovial membrane from OA patients, finding that approximately 50% of total live cells were PDPN⁺ fibroblasts and 28% were immune cells. Synovial fibroblasts expressed TLR family members 1 through 9 to varying degrees. Upon stimulation of TLR1/2, TLR4, TLR5, and TLR2/6, these fibroblasts upregulated inflammatory factors, including IL-6, IL-8, and G-CSF, as well as cartilage-degrading enzymes such as *MMP3* and *MMP10*. Interestingly, TLR1/2 stimulation did not affect the metabolic activity of synovial fibroblasts. However, when TLR1/2-prestimulated fibroblasts were cocultured with chondrocyte spheroids, they suppressed the expression of the cartilage-anabolic factor *COL2A1* and increased the expression of the cartilage-catabolic factors *MMP3* and *MMP10*, along with the inflammatory mediators IL-6, IL-8, and G-CSF. Consequently, chondrocyte spheroids exhibited reduced growth. These findings indicate that TLR1/2 pre-stimulation primes synovial fibroblasts to impair chondrocyte function.

In RA, synovial fibroblasts are known to express TLRs, particularly TLR2 and TLR4 [26]. Activation of TLR signaling in these cells triggers downstream pathways, such as NF-κB, leading to the production of pro-inflammatory mediators including IL-1β, TNF, and IL-6, as well as matrix-degrading enzymes such as MMPs that contribute to cartilage and bone destruction [40]. In the present study, we demonstrated that synovial fibroblasts from OA patients also express multiple TLRs. Similar to their counterparts in RA, OA synovial fibroblasts respond to TLR stimulation by upregulating inflammatory cytokines and cartilage-degrading enzymes. These findings suggest that synovial fibroblasts from RA and OA patients share common TLR-mediated response patterns, highlighting overlapping pro-inflammatory mechanisms in both diseases.

TLR signaling plays a significant role in OA development. In a high-fat diet–accelerated surgical OA model, knockout of *Tlr2*, but not *Tlr4*, mitigated OA severity, highlighting a pathogenic role of *Tlr2* in low-grade inflammation–driven OA in mice [41]. Low-grade inflammation in OA has traditionally been attributed to innate immune cells, such as synovial macrophages and neutrophils, with contributions from the complement system [42–44]. Our earlier work expanded this view by demonstrating that human chondrocytes themselves actively contribute to the low-grade innate immune response via TLRs [5]. Upon TLR activation, chondrocytes secreted a broad range of inflammatory mediators—including TNF, IFNγ, IL-6, IL-8, and G-CSF—which can either directly induce cartilage breakdown by activating chondrocytes or recruit macrophages and neutrophils to the synovial membrane, leading to synovitis.

The present study extends this concept further by identifying synovial fibroblasts from OA patients as additional innate immune players. These fibroblasts express a broad spectrum of TLR family members and respond to TLR stimulation with robust production of inflammatory cytokines and cartilage-degrading enzymes. Using a co-culture system designed to mimic the in vivo anatomical relationship between synovial membrane and cartilage, we demonstrated the deleterious effects of TLR1/2-prestimulated fibroblasts on chondrocyte function. Together, the interactive inflammatory cascades between chondrocytes in cartilage and fibroblasts in the synovial membrane likely form a positive feedback loop that promotes OA progression.

In response to TLR stimulation, we recorded that chondrocytes and synovial fibroblasts exhibited both shared and distinct molecular responses. Both cell types upregulated the expression of inflammatory cytokines, including IL-6, IL-8, and G-CSF, as well as the cartilage-degrading enzyme *MMP3*. In addition, both showed reduced ROS accumulation. However, synovial fibroblasts neither upregulated *NOS2* expression nor increased NO production, in marked contrast to the strong induction of both *NOS2* and NO observed in chondrocytes. Consistently, synovial fibroblasts did not exhibit impaired mitochondrial respiratory capacity, whereas chondrocytes showed pronounced mitochondrial dysfunction in response to TLR1/2 stimulation. The absence of excessive NO production and the preserved mitochondrial respiration in synovial fibroblasts further support the critical role of NO in regulating mitochondrial function, as previously demonstrated in chondrocytes [5].

Articular chondrocytes in cartilage and synovial fibroblasts in the synovial membrane are physically separated but communicate through the shared synovial fluid. To model this interaction, we established a Trans-well-based co-culture system in which synovial fibroblasts were seeded as a monolayer in the lower chamber, while chondrocytes were cultured as spheroids in the upper insert. Using this system, we found that TLR1/2-prestimulated OA synovial fibroblasts suppressed the expression of cartilage-anabolic markers such as *COL2A1*, while inducing the cartilage-catabolic enzymes *MMP3* and *MMP13* in chondrocytes. In addition, co-cultured chondrocytes exhibited increased expression of inflammatory genes, including IL-6, IL-8, and G-CSF. Consistent with these molecular changes, chondrocyte spheroids displayed reduced growth when co-cultured with TLR1/2-prestimulated fibroblasts, indicating impaired cartilage-forming capacity.

Despite these insights, this experimental design has inherent limitations. In the synovial membrane, several other cell types, including macrophages, neutrophils, and mast cells, also express TLRs and actively participate in inflammatory responses [45–47]. To more comprehensively assess the impact of TLR agonists within the synovial cavity on chondrocyte function, future studies could evaluate the responses of these additional TLR-expressing cells and their interactions with chondrocytes. Incorporating these immune cell populations, both individually and in combination with synovial fibroblasts, into co-culture systems could provide a more comprehensive model of the joint microenvironment.

Our findings also have important translational implications for the treatment of OA. Given the prominent role of TLR signaling in driving inflammatory and cartilage-catabolic responses in synovial fibroblasts and chondrocytes, therapeutic strategies targeting this pathway—currently under evaluation in RA [48]—may represent promising interventions for OA. These approaches include the use of blocking antibodies that prevent ligand-mediated TLR activation, small-molecule inhibitors that disrupt downstream signaling pathways, such as Auranofin [49] and CPG-52364 [50], and agents targeting key adaptor proteins including MYD88 and MAL. By attenuating TLR-mediated inflammatory signaling, these therapies have the potential to limit cytokine production, matrix degradation, and cartilage damage in OA. Notably, local intra-articular delivery of TLR-targeting agents may maximize therapeutic efficacy while minimizing systemic side effects. Collectively, our results provide a rationale for further preclinical and clinical evaluation of TLR-focused therapies as disease-modifying strategies in OA.

## Supporting information

Supplemental 1-6, and will be used for the link to the file on the preprint site.

Supplemental methods, and will be used for the link to the file on the preprint site.

## Acknowledgments

We thank Vivien Holecska, Katrin Lehmann, and Maria Dzamukova for experimental assistance, DRFZ FACS core facility, especially Jenny Kirsch and Kerstin Heinrich for cell sorting, and Philippe Saikali, Frank Zaucke, Antigoni Triantafyllopoulou, and Tonia Vincent for scientific discussions.

## Author Contributions

PS and ML designed the research. YD, AG, ND, XL, XL, YL, PW, and SW performed the experiments and analysed the data. KL, and LJG, PD, FH, and MFM conducted the RNA-sequencing and the corresponding computational data analysis. SO, LL, LP, TM, SD, TW, and CP coordinated patient sample collection. PS wrote the manuscript. ML, EL, and GK revised the manuscript. ML oversaw the research program, and received funding.

## Declaration of generative AI in scientific writing

We didn‘t use generative AI or AI-assisted technologies in preparing this work.

