## Supplemental 1-6, and will be used for the link to the file on the preprint site. for "TLR-mediated activation of synovial fibroblasts from osteoarthritis patients promotes chondrocyte dysfunction"

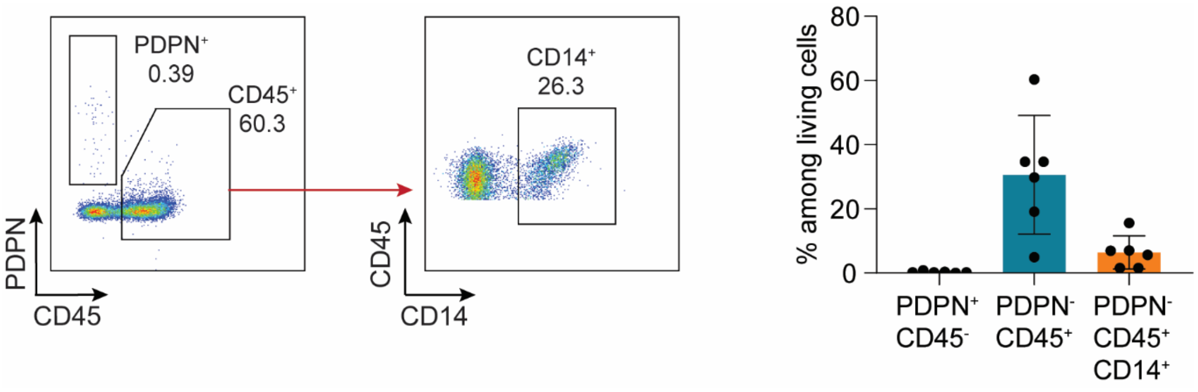


**Fig. S1. Synovial uid of OA patients is virtually devoid of -broblasts.** Synovial fluid of OA patients was digested with hyaluronidase (50 µg/ml) at 37°C for 30 minutes. Single-cell suspensions were analyzed by flow cytometry. Left, representative FACS plots. Right, quantification of fibroblasts (PDPN$^{+}$CD45$^{-}$), total immune cells (PDPN$^{-}$CD45$^{+}$), and monocytes/macrophages (CD45$^{+}$CD14$^{+}$), shown as percentages (n = 6, mean ± SD).

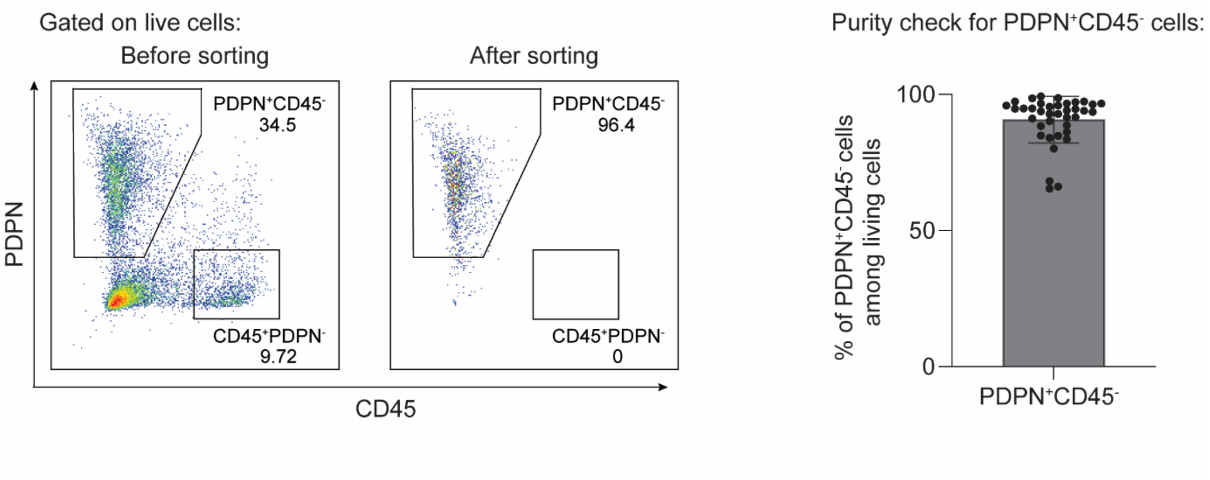


**Fig. S2. Fibroblast sorting from synovial membrane and purity check.** Synovial membrane tissues were collected from OA patients immediately after surgery. Isolated synovial cells were stained with anti-PDPN and anti-CD45. PDPN$^{+}$CD45$^{-}$ synovial fibroblasts were isolated by fluorescence-activated cell sorting (FACS) and immediately subjected to purity check. Left, representative FACS plots. Right, purity check for fibroblasts (PDPN$^{+}$CD45$^{-}$) (n = 37, mean ± SEM).

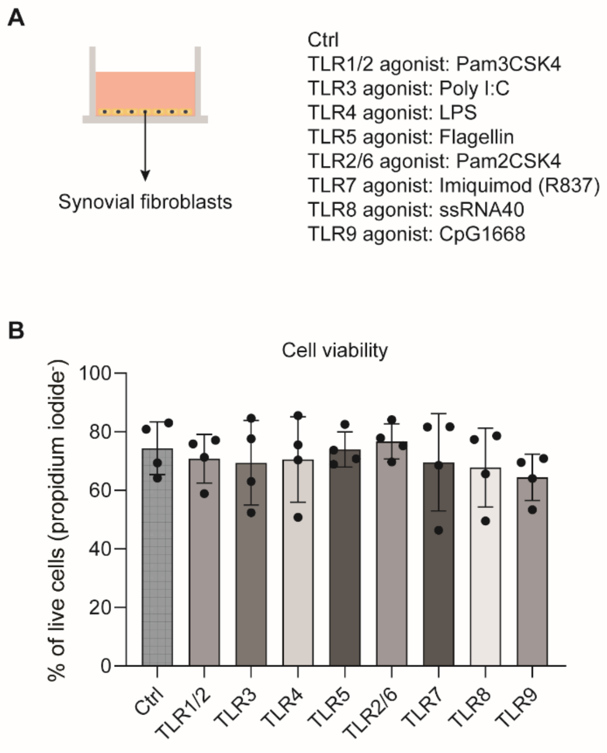


**Fig. S3. TLR stimulation setup and viability of purified synovial fibroblasts.** (**A**) Freshly purified synovial fibroblasts were stimulated with agonists targeting TLR1 to TLR9. (**B**) Four days later, fibroblast cells were harvested and stained with propidium iodide to determine the viability (n=4, mean ± SD).

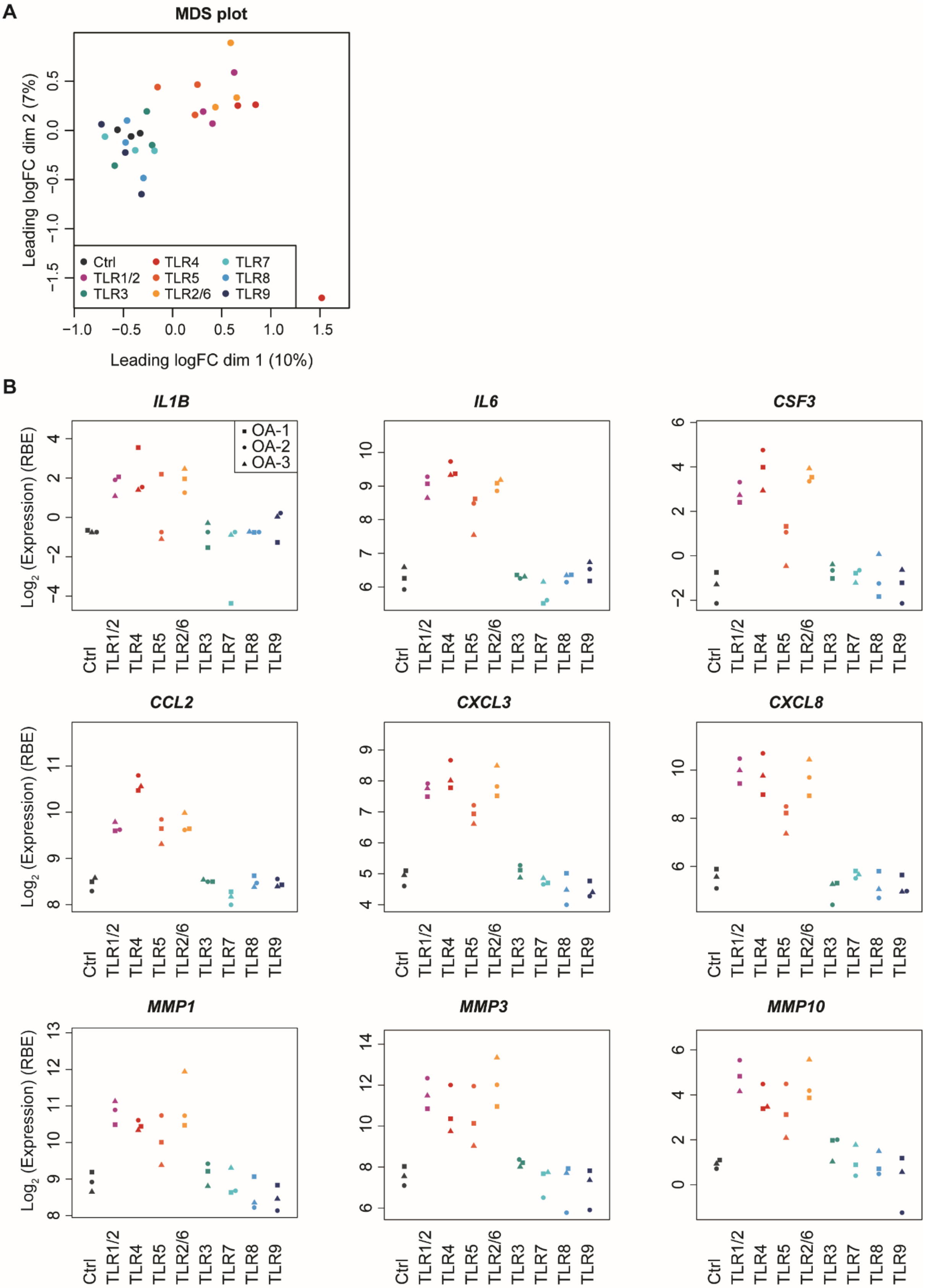

A
MDS plot
Leading logFC dim 2 (7%)
Leading logFC dim 1 (10%)
Ctrl
TLR1/2
TLR3
TLR4
TLR5
TLR2/6
TLR7
TLR8
TLR9
B
IL1B
IL6
CSF3
CCL2
CXCL3
CXCL8
MMP1
MMP3
MMP10
OA-1
OA-2
OA-3
Log2 (Expression) (RBE)

**Fig. S4. Synovial fibroblasts respond differently to distinct TLR stimulations.** TLR-stimulated fibroblast cells were harvested and subjected to bulk RNA-sequencing, using unstimulated cells as a control (Ctrl) (n=3). (**A**) Multidimensional Scaling (MDS) plot to visualize the level of similarity among these different treatments. The donor effect was first removed from the expression data. (**B**) The expression of prototypical OA inflammatory factors and cartilage-degrading enzymes in TLR1- to TLR9-stimulated fibroblasts, as well as unstimulated control cells.

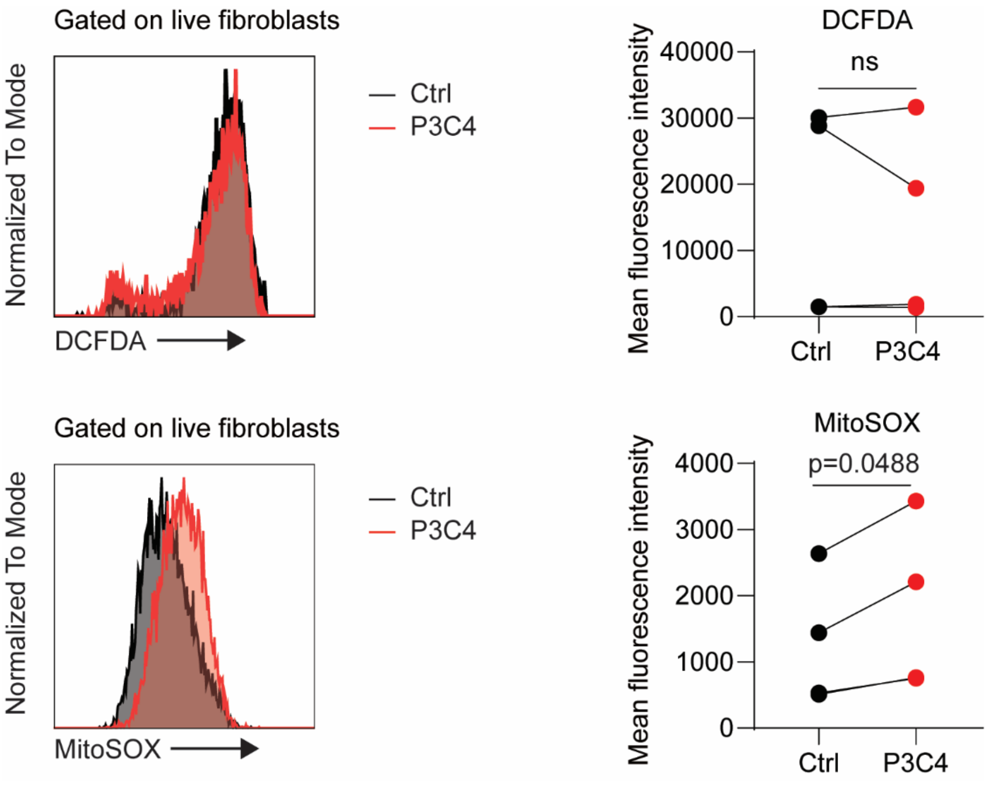


**Fig. S5. TLR1/2-stimulation slightly increases mitochondrial ROS production of synovial fibroblasts**. Freshly purified synovial fibroblasts were stimulated with Pam3CSK4 (P3C4), the agonist of TLR1/2, or were left unstimulated as a control (Ctrl). Four days later, cells were harvested and subjected to DCFDA and MitoSOX staining to detect the total intracellular ROS and mitochondrial ROS, respectively (n=3–4). Left: representative graphs; right: quantification. Data were compared using paired two tailed t test.

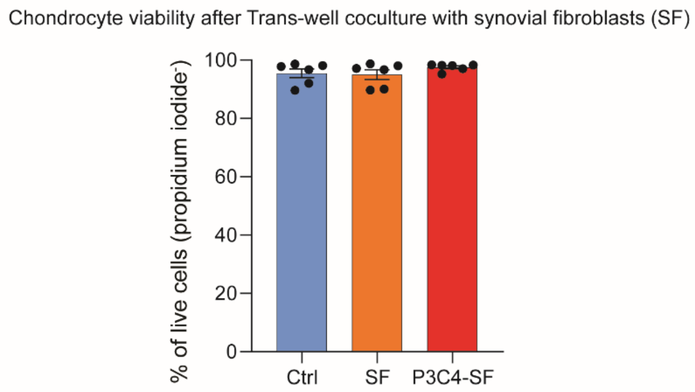


**Fig. S6. Chondrocyte viability after Trans-well coculture with synovial fibroblasts.** After a 4-day co-culture described in figure 4, chondrocyte spheroids were dissociated with collagenase II to generate single chondrocytes, which were subsequently stained with propidium iodide for viability determination. Ctrl: chondrocytes alone; SF: synovial fibroblast–stimulated chondrocytes; P3C4-SF: Pam3CSK4-prestimulated synovial fibroblast–stimulated chondrocytes.
