## Supplemental methods, and will be used for the link to the file on the preprint site. for "TLR-mediated activation of synovial fibroblasts from osteoarthritis patients promotes chondrocyte dysfunction"

### Materials and Methods

#### Synovial fluid cell isolation

Synovial fluid samples without visible blood contamination were digested with hyaluronidase (50 µg/ml; Sigma, H3506) at 37°C for 30 minutes. The digested fluid was filtered through 70 µm MACS SmartStrainers and cells were washed three times before downstream analyses.

#### Chondrocyte isolation and spheroid culture

Cartilage samples were washed three times with DPBS (Gibco) and finely minced with scalpels. The tissue fragments were digested with collagenase II (1 mg/ml) for 16 hours. The resulting suspension was filtered through a 70 µm MACS SmartStrainer (Miltenyi) to obtain single chondrocytes, which were then washed and resuspended in culture medium. To generate spheroids, chondrocytes were resuspended in serum-free DMEM-High Glucose medium (Sigma Aldrich) supplemented with 0.1 µM dexamethasone (Sigma Aldrich; D2915), 40 µg/ml L-proline (Sigma Aldrich; P5607), 6.25 µg/ml insulin-transferrin-sodium selenite supplement (Sigma Aldrich; I1884), 0.1 mg/ml sodium pyruvate (AppliChem; A4859), 1.25 mg/ml bovine serum albumin (Sigma Aldrich; A9418), 1% penicillin-streptomycin (Gibco; 15140-122), 50 µg/ml 2-phospho-L-ascorbic acid trisodium salt (Sigma Aldrich; A8960), 5.35 µg/ml linoleic acid (Sigma Aldrich; L1012), and 10 ng/ml TGF-β1 (Peprotech; 100-21-5). For spheroid formation, $2.5 \times 10^5$ cells were transferred into 15 ml Falcon tubes and centrifuged at 500 × g for 5 minutes. After carefully removing the supernatant, 500 µl of fresh spheroid medium was added without disturbing the cell pellet. Cultures were maintained in a hypoxic incubator (4% $O_2$, 5% $CO_2$, 37°C), and medium was renewed every three days using a hypoxic chamber workstation (BioSpherix X3, Xvivo system).

#### Trans-well coculture

Isolated synovial fibroblasts were stimulated with or without the TLR2 agonist Pam3CSK4 (2 µg/ml) for 3.5 days in a 24-well plate. After stimulation, fibroblasts were harvested, washed twice, resuspended in chondrocyte spheroid culture medium (survival was comparable to standard fibroblast culture medium), counted, and seeded into a fresh 24-well plate at the same cell density. A 0.4 µm Trans-well insert (Millipore) was then placed into each well, and 0.5 ml of culture medium was added to the upper chamber. Chondrocyte spheroids derived from the same patients were transferred onto the inserts, allowing fibroblasts in the lower chamber and chondrocyte spheroids in the upper chamber to share medium while remaining physically separated. Cocultures were maintained in a hypoxic incubator (4% $O_2$, 5% $CO_2$, 37°C) with medium refreshed every 3.5 days. Supernatants were collected at each medium change and stored at –20°C for further analyses.

**Flow Cytometric Analysis**

Cell suspensions were first treated with Human TruStain FcX (Fc receptor blocking solution, Biolegend, 1:20) for 10 minutes at 4°C to prevent nonspecific binding. Cells were then washed with PBS containing 1% BSA and stained with the antibody mixture for 45 minutes at 4°C. For intracellular staining, cells were fixed with 2% paraformaldehyde (PFA) for 10 minutes at room temperature, washed with 0.05% saponin, and incubated with antibodies diluted in 0.05% saponin for 30 minutes at 4°C. The following antibodies were used: Anti-hPDPN (Invitrogen; 12-9381-42), Anti-hCD90 (Biolegend; 328116), Anti-hCD45 (Biolegend; 368532), Anti-hCD14 (Biolegend; 325615), Anti-hCD3 (Biolegend; 317418), Anti-hCD8 (Biolegend; 301035), Anti-hCD19 (Biolegend; 302258), Anti-hTLR1 (Abcam; ab59702), Anti-hTLR2 (Abcam; ab13553), Anti-hTLR3 (Abcam; ab45093), Anti-hTLR4 (Abcam; ab8378), Anti-hTLR5 (R&D Systems; FAB6704G), Anti-hTLR6 (Abcam; ab72362), Anti-hTLR7 (R&D Systems; IC5875P), Anti-hTLR8 (Biolegend; 395507), Anti-hTLR9 (Abcam; ab58864), Zombie Aqua Fixable Viability Kit (Biolegend; 423102), and Zombie NIR Fixable Viability Kit (Biolegend; 423106). To acquire apoptotic status, cells were stained with PI and Annexin V. Stained cells were acquired on a BD FACS Canto II flow cytometer and analyzed using FlowJo software (version 10.7.1).

**TLR stimulation**

Isolated synovial fibroblasts were stimulated with 2 µg/ml of Pam3CSK4, PolyI:C, LPS, Flagellin, Pam2CSK4, Imiquimod, ssRNA, or CpG (InvivoGen), which act as agonists of TLR1/2, 3, 4, 5, 2/6, 7, 8, and 9, respectively.

**mRNA isolation and quantitative reverse transcription PCR**

For synovial fibroblasts, cells were harvested, washed, and lysed, and total RNA was extracted using the NucleoSpin RNA XS Micro Kit (MACHEREY-NAGEL). Extracted RNA was used for either reverse transcription followed by qPCR or for bulk RNA sequencing. Chondrocyte spheroids were transferred to 1 ml Lysis/Binding Buffer (µMACS mRNA Isolation Kit, Miltenyi Biotec) and mechanically disrupted using a gentleMACS device with M tubes (Miltenyi Biotec). mRNA was then isolated using Oligo (dT) magnetic beads (µMACS mRNA Isolation Kit, Miltenyi Biotec) following manufacturer's instructions. cDNA was reverse-transcribed from isolated mRNA using TaqMan reverse transcription reagents (Thermo Fisher Scientific). Expression of target genes was quantified by qPCR using Fast SYBR Green Master mix reagents and Quant Studio 7 devices (Thermo Fisher Scientific). Relative expression levels were calculated using the ΔΔCt method with *ACTB* as the housekeeping control. Primers (forward and reverse) were obtained from Eurofins Genomics: *ACTB* FP: 5'-CACCCAGCACAATGAAGATCAAGA-3', *ACTB* RP: 5'-CCAGTTTTTAAATCCTGAGTCAAGC-3'; *COL2*A1 FP: 5'-GGAATTCGGTGTGGACATAGG-3', *COL2A1* RP: 5'-ACTTGGGTCCTTTGGGTTTG-3'; *ACAN* FP: 5'-GAATGGGAACCAGCCTATACC-3', *ACAN* RP: 5'-TCTGTACTTTCCTCTGTTGCTG-3'; *MMP1* FP: 5′-AAAATTACACGCCAGATTTGCC-3′, *MMP1* RP: 5′-GGTGTGACATTACTCCAGAGTTG-3′; *MMP3* FP: 5'-TTTTGGCCATCTCTTCCTTCA-3', *MMP3* RP: 5'-TGTGGATGCCTCTTGGGTATC-3'; *MMP13* FP: 5′-TCCTGATGTGGGTGAATACAATG-3′, *MMP13* RP: 5′-GCCATCGTGAAGTCTGGTAAAAT-3′; *ADAMTS1* FP: 5′-GGACAGGTGCAAGCTCATCTG-3′, *ADAMTS1* RP: 5′-TCTACAACCTTGGGCTGCAAA-3′; *ADAMTS4* FP: 5′-GAGGAGGAGATCGTGTTTCCA-3′, *ADAMTS4* RP: 5′-CCAGCTCTAGTAGCAGCGTC-3′; *ADAMTS5* FP: 5'-GCTCACGAAATCGGACATTTACTT-3', *ADAMTS5* RP: 5'-ACCAAGGTCTCTTCACAGAATTTG-3'; *IL6* FP: 5'-ATGAACTC CTTCTCCACAAGC-3', *IL6* RP: 5'-GTTTTCTGCCAGTGCCTCTTTG-3'; *CXCL8* FP: 5'-

GGCACAAACTTTCAGAGACAGCAG-3’, *CXCL8* RP: 5’-GTTTCTTCCTGGCTCTTGTCCTAG-3’; *CSF3* FP: 5’-TGAGTGTGCCACCTACAAGC-3’, *CSF3* RP: 5’-GACACCTCCAGGAAGCTCTG-3’; *NOS2* FP: 5’-GTTCTCAAGGCACAGGTCTC-3’, *NOS2* RP: 5’-GCAGGTCACTTATGTCACTTATC-3’.

**Synovial fibroblast TaqMan qPCR**

After FACS sorting, $PDPN^+$ synovial fibroblasts were lysed with TRIzol buffer, and total RNA was extracted using the RNeasy Mini Kit (Qiagen; 217004). cDNA was transcribed using TaqMan reverse transcription reagents (Thermo Fisher Scientific; N8080234). TaqMan qPCR were performed in a Quant Studio 7 device utilizing TaqMan™ Fast Advanced Master Mix (Thermo Fisher; 4444556) in combination with the following TaqMan Gene Expression assays: Hs00413978_m1 (*TLR1*), Hs00610101_m1 (*TLR2*), Hs01551078_m1 (*TLR3*), Hs00152939_m1 (*TLR4*), Hs00152825_m1 (*TLR5*), Hs00271977_s1 (*TLR6*), Hs00152971_m1 (*TLR7*), Hs00152972_m1 (*TLR8*), Hs00152973_m1 (*TLR9*), Hs01935337_s1 (*TLR10*), Hs01573837_g1 (*MYD88*), Hs01090712_m1 (*TICAM1*), Hs01019083_m1 (*VDAC1*), Hs03023943_g1 (*ACTB*).

**RNA-sequencing analysis**

RNA integrity was assessed using a Fragment analyzer (Agilent) and cDNA libraries were generated for samples with high RNA integrity (RQN > 8), using the Smart-Seq mRNA LP Kit (Clontech) with up to 10 ng of RNA according to manufacturer’s instructions. Paired-end sequencing (2x111 bp) of cDNA libraries was performed on an Illumina NextSeq 2000 device. Obtained reads were mapped to the hg19 genome (annotation releases: GRCh37.p13) using Tophat2 (25) and Bowtie2 (26) with very-sensitive settings. Read counts were determined with featureCounts (27).

RNA-seq data were analyzed in R (v4.4.0) (https://www.R-project.org/). Read alignment was performed using the align function of the Rsubread (v2.18.0) package (28). An index was built using the Ensembl Homo sapiens GRCh38 primary assembly genome file, and FASTQ files were aligned with default settings. Unstranded gene-level counts were then obtained using the featureCounts function and the Ensembl Homo sapiens GRCh38 GTF annotation file (v115).

Differential gene expression analyses were performed using the edgeR (v4.2.0) (29) and limma (v3.60.0) packages (30). A DGEList object was created from the featureCounts output and Ensembl gene IDs were annotated using the biomaRt package (31, 32). A design matrix was made incorporating sample treatment group and donor. Lowly expressed genes were removed using the filterByExpr function and normalization factors were calculated using the TMM method (33).

For plotting gene expression, log2 counts per million (CPM) expression values were calculated using the edgeR cpm function. This was performed after filtering and TMM normalization (except when plotting TLR gene expression). The donor effect was then subtracted using the removeBatchEffect function. A multidimensional scaling (MDS) plot was created with the plotMDS function using the top 500 most variable genes for each sample pairing. Relative log2 CPM expression values were calculated for each gene by subtracting the average log2 CPM across all control samples, with expression scales truncated at ±4. Heat maps were made using the pheatmap package (v1.0.12).

Counts were transformed using the voom method (34), and a linear model was fit using the edgeR voomLmFit function and the design matrix. Groups were compared using the contrasts.fit function and moderated t-statistics were calculated using eBayes (35). Differentially expressed genes were defined as those with a false discovery rate (FDR)-adjusted p-value < 0.05. For each comparison, genes were ranked by confident effect size (confect) using the limma_confects function from the topconfects package (v1.20.0) (36). Differentially expressed genes were shown in mean-difference (MD) plots, highlighting genes that were significant at various confect thresholds.

The Reactome (37) gene set collection was obtained from the Broad Institute Molecular Signature Database (38, 39), via the msigdbr package (v7.5.1). Gene set testing was performed using the cameraPR function (40) from the limma package on the moderated t-statistics. The top ten most significant gene sets across comparisons of interest were presented as a heat map showing the average log2 fold changes of genes in each set (rows) for the different comparisons (columns), with the scale truncated at ±2. Gene sets were manually curated into the following groups: extracellular matrix homeostasis, inflammation, GPCR signaling, antigen presentation and metal transporter. The significance of each gene set was indicated by the text (−log10 FDR-adjusted p-value threshold; an asterisk denotes adjusted $p < 0.05$, 2 denotes $p < 0.01$, 3 denotes $p < 0.001$ and so on).

**Bio-plex analysis**

Culture supernatants from control or TLR-stimulated conditions were collected and subjected to Bio-plex assay (Bio-Rad) to quantify the concentration of IL-6, IL-8, and G-CSF.

**Nitric oxide quantification by modified Griess reaction**

Culture supernatants from control or TLR-stimulated conditions were collected and nitrite/nitrate oxidized from NO was measured spectrophotometrically after addition of Griess reagent (Sigma–Aldrich; G-4410 and InvivoGen) according to manufacturer´s instructions.

**MitoSpy, TMRM, MitoSox, and DCFDA staining**

Harvested cells were washed and incubated for 30 minutes at 37°C with MitoSpy (BioLegend), TMRM, MitoSOX (Thermo Fisher Scientific), and DCFDA (Abcam; ab113851), diluted in pre-warmed DMEM-High Glucose medium. Following staining, LIVE/DEAD Fixable Near-IR Dead Cell Stain (Thermo Fisher Scientific) was added to identify dead cells. Samples were acquired on a BD FACSCanto II flow cytometer and analyzed using FlowJo software (version 10.7.1).

**Mito Stress Test seahorse assay**

After 4-day stimulation, control and TLR-stimulated fibroblasts were reseeded in Agilent Seahorse XFe96 Microplate wells, which were precoated with poly-D-lysine and contained prewarmed assay buffer. OCR and ECAR measurements were performed every 5 min prior to and after sequential addition of oligomycin, FCCP or Rotenone/Antimycin A. Data were analyzed using Wave (Agilent).

**Immunofluorescence Staining of Synovial Membrane**

Fresh synovial membrane samples were immediately fixed in 4% PFA at 4°C for 24 hours, washed three times with cold PBS, and sequentially immersed in 10%, 20%, and 30% sucrose solutions at 4°C for 24 hours each. Samples were embedded in SCEM medium (Section Lab) and frozen using OCT medium (Tissue-Tek). Sections were cut at 6 µm thickness on a cryotome. Hydrophobic barriers were drawn around sections using a PAP pen and sections were rehydrated with PBS for 5 minutes at room temperature (RT). Sections were permeabilized with 0.3% Triton X-100 and incubated overnight at 4°C with primary antibodies: Podoplanin Monoclonal Antibody (Invitrogen; 14-9381-82) and Human TLR2 Antibody (R&D Systems; AF2616). Sections were then brought to RT for 15 minutes, washed three times with PBS-0.1% Tween 20, and incubated for 1 hour at RT in the dark with secondary antibodies (DyLight 550, Invitrogen; SA5-10027, and Alexa Fluor® 594, Life Technologies; A-11058). Sections were washed three times with PBS and counterstained with DAPI (1:250) for 10 minutes at RT in the dark. After final washes, mounting medium and coverslips were applied, and sections were allowed to dry in the dark for 30 minutes. Confocal images were acquired using a Zeiss LSM880 microscope.

**Statistics**

Statistical analysis was performed using GraphPad Prism (v5.02 and v7). Data were first examined for normality. If a normal distribution was found, significance was determined using paired or unpaired two-tailed t-tests for two-group comparisons and a one-way ANOVA was used for multiple group comparisons. In case of non-normally distributed groups, comparisons were performed using nonparametric tests with corresponding corrections (paired t-test: Wilcoxon correction; unpaired t-test: Mann-Whitney U test; One-Way ANOVA: Friedman test). For comparison of two groups in kinetic analyses, a Two-way ANOVA was used.

**Study approval**

The study was approved by the Ethics Committee of Charité – Universitätsmedizin Berlin (EA4/022/21).
